# Parallel Framework for Inferring Genome Scale Gene Regulatory Networks

**DOI:** 10.1101/2021.07.11.451988

**Authors:** Softya Sebastian, Swarup Roy

## Abstract

Genome-scale network inference is essential to understand comprehensive interaction patterns. Current methods are limited to the reconstruction of small to moderate-size networks. The most obvious alternative is to propose a novel method or alter existing methods that may leverage parallel computing paradigms. Very few attempts also have been made to re-engineer existing methods by executing selective iterative steps concurrently. We propose a generic framework in this paper that leverages parallel computing without re-engineering the original methods. The proposed framework uses state-of-the-art methods as a black box to infer sub-networks of the segmented data matrix. A simple merger was designed based on *preferential attachment* to generate the global network by merging the sub-networks.

Fifteen (15) inference methods were considered for experimentation. Qualitative and speedup analysis was carried out using DREAM challenge networks. The proposed framework was implemented on all the 15 inference methods using large expression matrices. The results were auspicious as we could infer large networks in reasonable time without compromising the qualitative aspects of the original (serial) algorithm.

CLR, the top performer, was then used to infer the network from the expression profiles of an Alzheimer’s disease (AD) affected mouse model consisting of 45,101 genes. We have also highlighted few hub genes from the network that are functionally related to various diseases.

## 1. Introduction

The complicated interactions that transpire between the biological molecules such as gene, mRNA and protein are responsible for the disparate cellular functions that occur within an organism. These relations or interactions are often represented using a biomolecular interaction network or graph in which each node is a biomolecule and the edge represents the interaction or the association between two such biomolecules or a complex combination of biomolecules [1, 2]. Biomolecular networks include gene regulatory networks (gene-gene interactions), transcription regulatory networks (transcription factor-gene interactions), signalling networks (integrated interactions among molecules), protein interaction networks (protein-protein interactions), metabolic networks (enzyme-substrate interactions), and hybrid networks. All these networks imperatively exist in all cell systems to fulfil their fundamental and essential roles in not only procreating but also sustaining life.

System and molecular biological research over the years have shifted the focus from analyzing individual components to investigating biomolecular networks. High-throughput experimental methods have made it easier to study biomolecular networks by providing enormous data about interactions, networks, functional modules [3, 4], and pathways. Gene Regulatory Networks (GRNs) offer a lot of insight into the mechanisms of the complex cellular systems and hence the inference of GRN is indispensable in the quest to comprehend genes, their functions as well as their relationship with other genes. Consequently, a plethora of methods for the inference of gene regulatory networks from gene expression profile has been proposed in the last few decades [5, 6, 7, 8, 9, 10, 11, 12, 13, 14, 15, 16, 17, 18]. A majority of the methods employ an expensive association measure to draw the possible edge between a pair of nodes (genes), making the large network inference a hard task. Hence, they are limited to handling just a few hundred genes. In reality, genome-scale network inference is the ultimate need in network biology [1]. The task of inference of large scale gene regulatory networks is a complex optimisation problem that draws in an enormous amount of gene expression data from thousands of genes with multiple time-series expressions or samples. Handling this massive data in addition to its complexity makes the inference of large scale gene regulatory networks a computationally expensive task.

In addition to the qualitative aspect of the inference methods, the scalability of the methods is equally important. Very little is known about the scalability of such methods in reconstructing genome-scale networks. The development of alternative concurrent computing platforms may massively accelerate the compute-intensive inference task. Very few attempts have been made towards the same. One common approach is to pick up a comparatively good inference method and parallelly run a few of its steps with some modifications and obtain the result. Instead of proposing a new method or extending existing methods, one alternative is to utilise the available inference methods by empowering them to infer relatively larger GRNs parallelly without altering their original structure. The contributions of this paper are listed next.

- A simple yet effective generic parallel framework that enables serial inference methods to infer large networks parallelly without compromising on their quality has been proposed. This framework virtually makes any number of existing methods capable of inferring large scale networks without the need to re-engineer them.
- Fifteen (15) candidate state-of-the-art inference methods were categorised based on the core methodologies that they adopted. The scalability analysis of these 15 inference algorithms together with their qualitative performance was assessed.
- The candidate inference methods were ranked based on their qualitative and quantitative performances.
- The proposed generic parallel framework was implemented on the candidate inference methods and the effectiveness of the framework was analysed and reported.
- The top-ranked inference method was further employed to infer the genome scale network of Alzheimer’s disease (AD) affected *mouse* genes which was then analysed for prioritising possible disease responsible genes. We were successful in reconstructing a network consisting of 45,101 nodes (genes) in a reasonable amount of time. To the best of our knowledge, no prior work has experimented with such a large scale network and downstream network analysis.

## 2. Traditional Gene Regulatory Network Inference Methods

In this section, the popular candidate GRN inference methods selected for the scalability analysis as well as for demonstrating the efficacy of the generic parallel framework proposed in this paper are briefly introduced.

The huge amount of transcriptome data that are regularly generated and updated require fast and accurate computational tools for processing and analysis to be productive. In this regard, many GRN inference methods have been proposed and implemented in the past two decades. There are different approaches to reconstructing gene regulatory networks and different algorithms have been proposed based on each of these approaches. Although the methods have been presented with the hope of delivering promising results, very little is mentioned about their applicability especially in terms of their scalability with varying expression matrix size. Herein, 15 inference algorithms were shortlisted on the basis of their popularity, and the availability of their original implementation (codes). The different algorithms have been categorised based on the computational models they adopt to infer the network.

### 2.1. Mutual Information-based Inference

Mutual information-based GRN inference is one of the most popular and widely used approaches in inferring GRNs. Mutual information score between the expression levels of two genes represents the degree of dependence of the two genes. Score zero indicates observations are independent, otherwise it is the entropy of the marginal distribution. Mutual information networks are undirected since the mutual information scores are symmetric. A total of 6 MI-based inference methods were considered because of their overwhelming popularity as well as easy availability of code and applicability.

**RelNet** (Relevance Network) [5] infers an edge between two genes in a network only if the mutual information of the two genes is larger than a certain threshold. All pairwise interactions are assessed herein and since the method does not eliminate the indirect interactions between genes, there is always a consequence that two genes may be shown as interacting, even though it may not be the case. **ARACNE** (Algorithm for the Reconstruction of Accurate Cellular Networks) [7] seeks to overcome this limitation using the Data Processing Inequality (DPI). It first identifies candidate interactions by estimating pairwise mutual information, filters them and then the indirect candidate interactions are removed using DPI. Then came **CLR** (Context Likelihood of Relatedness) algorithm [6] that applies an adaptive background correction step to eliminate false correlations and indirect influences to overcome the limitations of relevance networks. The mutual information is computed first and then CLR calculates the statistical likelihood of each mutual information value within its network context.

**C3NET** (Conservative Causal Core NETwork) algorithm [10] identifies direct physical gene interactions from expression data by identifying a significant maximum mutual information network such that two genes are connected only if their shared significant mutual information value is at least maximal with respect to all the other genes. **C3MTC** [11] is an improvised modification of C3NET [10] that uses an additional multiple-testing procedure to control the type one error for a more efficient implementation. **BC3NET** [12] generates an ensemble of independent bootstrap datasets by sampling using a non-parametric bootstrap from one dataset and then for each generated data set in the ensemble, a network is inferred using C3NET. From this ensemble of networks, one weighted network is constructed by determining the statistical significance of the connection between gene pairs which results in the final binary undirected network.

### 2.2. Statistical Measures-based Inference

Approaches based on statistical measures reconstruct a network by computing a similarity score, say, correlation, for each pair of genes wherein an edge is inferred only if the score is greater than a certain threshold. The inferred network is undirected because these scores are symmetric. This approach requires a post-processing step to weed out the indirect regulatory relationships. Six inference methods based on this approach were identified.

**MutRank** [13] first ranks the correlation between every pair of genes using Pearson’s correlation. An weighted edge is then assigned between genes *i* and *j* based on the geometric mean of the scores obtained between gene i and j and vice versa. **PCOR** [14] calculates the partial correlations for each pair of genes to reconstruct the network. The key principle of correlation networks is that if two genes have highly-correlated expression patterns meaning that they are co-expressed, then they are assumed to participate together in a regulatory interaction [19]. **SPCOR** [14] calculates the semi-partial correlations of all pairs of two random genes to build the network. While partial correlation gives a measure of the relationship between two variables or genes after eliminating the effect of all the other genes, semi-partial correlation only eliminates the effect of a fraction of other genes. **GeneNet** [16] on the other hand infers the network by first converting an inferred correlation network into a partial correlation graph and then constructing a directed acyclic causal network out of the partial correlation network. Whereas **MINE** (Maximal Information-Based Nonparametric Exploration) [18] is a statistic that provides quantitative evaluations of five different aspects of the relationship between two variables, out of which one is Maximal Information Coefficient. It shows the relationship strength and can be interpreted as a correlation measure that is symmetric and ranges in [0,1], where it tends to 0 for statistically independent data while it approaches 1 for noiseless functional relationships.

**G1DBN** [17] is based on the concept of low order conditional dependence graph which is extended to Dynamic Bayesian Networks (DBN) [20]. DBNs are an easier extension of Bayesian Networks that is capable to infer interaction uncertainties among genes using probabilistic graphical models. DBN is usually used when we need to describe the qualitative nature of the dependencies between genes in a temporal process.

### 2.3. Feature Selection-based Inference

Feature selection based inference considers GRN inference as a feature selection problem where the goal is to find the true or candidate regulators for every gene. This approach aims at mapping the significant score between two genes in such a way that true regulations are always obtained by applying a threshold on the list of candidate regulations.

**MRNET** (Minimum Redundancy NETworks) [8] is inspired by the feature selection technique called the maximum relevance/minimum redundancy (MRMR) algorithm. MRNET formulates a series of input/output supervised gene selection procedures and then adopts the MRMR principle to perform the gene selection for each supervised gene selection procedure. The idea is to perform a series of supervised MRMR gene selection procedure for every gene by putting in the expression levels of all genes. The algorithm infers an edge between two genes only when either of them is a well-ranked predictor of the other. MRNET is based on a forward selection strategy and its limitation is that the quality of the selected subset would depend on the variable that was selected at first. **MRNETB** (Minimum Redundancy NETworks Backward) [9] is an improved version of MRNET that uses a backward selection strategy, followed by a sequential replacement which requires the same computational cost as forward selection strategy to overcome the limitation of MRNET.

The **GENIE3** (GEne Network Inference with Ensemble of trees) [15] uses a feature selection technique based on random forests [21] to solve the regression problem for each gene and obtain the network. The expression pattern of the target gene is predicted from the expression patterns of all transcription factors for each regression problems.

A summary of all the methods discussed above is presented in Table 1. Several methods have been proposed and updated over the years to infer gene regulatory networks. Despite this, researchers are still on the lookout for new and improvised methods. This is because the traditional methods are bogged down by many limitations. The most common one being that the scalability of these methods are unaccounted for since most methods do not use large datasets for their assessment. As all the methods need to find if there is any relation between every two genes, these methods tend to be computationally expensive. Therefore, serial execution of these methods for datasets consisting of a very large number of genes seems impossible using machines without large computational capacity. Therefore, it is quite obvious that parallel GRN inference methods became necessary which is why it led to many new kinds of research emerging with the concept of parallel computing. Some methods were redesigned from the existing traditional algorithms by parallelly executing some steps, while some others were a concoction of several different approaches to infer regulatory networks. These approaches were well received within the research community because of their ability to save time and infer large networks but to what extent is still not known. Now the situation is such that some methods have been updated to parallel versions while some have been abandoned altogether. What new researchers tend to ignore is that each method for inference had its own set of advantages and limitations over the other methods. Different inference methods are required to be used separately or as an ensemble. Therefore, we strongly felt that there is a need to propose a generic parallel framework such that any state-of-the-art inference algorithm can be chosen and be used to infer large networks, leveraging parallel computing and yet without altering the original algorithm.

**Table 1:** Synopsis of GRN Inference Methods

| Approach | Algorithm | Inferred Network Type | Package | Platform | Assessment based on (type of data) | Availability |
| --- | --- | --- | --- | --- | --- | --- |
| Mutual Information | RelNet (1999) | Undirected | MINET (2008) | R | Real (Saccharomyces cerevisia) | https://www.bioconductor.org/packages/minet/ |
|  | ARACNE (2006) | Undirected | MINET (2008) | R | Synthetic (Random network) & Real (Human B Cells) | https://www.bioconductor.org/packages/minet/ |
|  | CLR (2007) | Undirected | MINET (2008) | R | Real (Escherichia coli) | https://www.bioconductor.org/packages/minet/ |
|  | C3NET (2010) | Undirected | C3NET (2012) | R | Synthetic (E.coli) & Real (E.coli) | https://cran.r-project.org/package=c3net |
|  | C3MTC (2011) | Undirected | bc3net (2016) | R | Synthetic (Erdős-Rényi Networks) | https://CRAN.R-project.org/package=bc3net |
|  | BC3NET (2012) | Undirected | bc3net (2016) | R | Synthetic (Erdős-Rényi, E.coli) & Real (S. cerevisiae) | https://CRAN.R-project.org/package=bc3net |
| Statistical Measures | MutRank (2009) | Undirected | NetBenchmark (2020) | R | Synthetic (Arabidopsis, Human, Mouse, Rat) | https://www.bioconductor.org/packages/netbenchmark/ |
|  | PCOR (2015) | Undirected | PPCOR (2015) | R | - | https://CRAN.R-project.org/package=ppcor |
|  | SPCOR (2015) | Undirected | PPCOR (2015) | R | - | https://CRAN.R-project.org/package=ppcor |
|  | GeneNet (2007) | Directed | GeneNet (2020) | R | Real (Arabidopsis thaliana) | https://CRAN.R-project.org/package=GeneNet |
|  | G1DBN (2009) | Directed | G1DBN (2013) | R | Synthetic (Random Networks) & Real (S. cerevisiae) | https://CRAN.R-project.org/package=G1DBN |
|  | MINE (2011) | Undirected | MINERVA (2019) | R, C, Python, Matlab | Real (Yeast) | https://CRAN.R-project.org/package=minerva |
| Feature Selection | MRNET (2007) | Undirected | MINET (2008) | R | Synthetic (sRogers & SynTreN) | https://www.bioconductor.org/packages/minet/ |
|  | MRNETB (2010) | Undirected | MINET (2020) | R | Synthetic (DREAM4) | https://www.bioconductor.org/packages/minet/ |
|  | GENIE3 (2010) | Directed | GENIE3 (2020) | R, MatLab | Synthetic (DREAM4) & Real (E.coli) | https://bioconductor.org/packages/devel/bioc/html/GENIE3.html |

## 3. Materials and Method

In this section, we discuss the datasets used throughout this work, the proposed generic parallel framework, the candidate serial inference methods, the execution platform for implementation as well as the parameters used for assessment.

### 3.1. Datasets

The scalability of the inference methods together with their prediction quality can be assessed only if datasets not only of varying sizes but also with the underlying true network are available. While real networks come with the desired number of genes, unfortunately, this area lacks true gene expression networks with corresponding expression profiles for the set of target genes. Hence, several approaches report the performance based on synthetic datasets that provide the underlying true network along with the expression data but with limited network size. In order to have the best of both worlds, we decided to perform scalability analysis on both real as well as synthetic datasets so as to effectively show the trend of the inference methods with increasing network size. But since the true networks were only available for synthetic networks, the qualitative assessment was performed using the results of only the synthetic networks.

#### 3.1.1. Real Datasets

The publicly available expression profile of the triple transgenic mouse model of Alzheimer’s Disease (3xTg-AD) with the GEO Accession # GSE32536 contributed by [22] was downloaded from National Center for Biotechnology Information (NCBI)^1^ to be employed as the real dataset. Searcy et.al [22] in their work experimented on 10-month-old triple transgenic mice with established amyloid-*β* (A*β*) deposition and pathology still ongoing (AD affected), that were treated with Thiazolidinediones (TZD) - pioglitazone (PIO) for four months. TZD-PIO is a drug that is used clinically for treating type 2 diabetes and is shown to re-establish insulin sensitivity, improve lipid profiles and reduce inflammation. Later it was also found to be beneficial in treating AD by ameliorating cognitive decline that occurs early in the disease and hence was used in the study. After the four months of treatment, they found that PIO treatment improved cognitive impairment, reduced A*β* and tau staining and altered expression for several pharmacologically-relevant gene targets. The dataset consisted of an expression profile of 45,101 genes. Subsets of required network size (10k, 15k and 20k genes) were extracted from this dataset to test the ability of the methods to handle large networks. However, due to lack of an underlying true or gold network for the dataset, the prediction accuracy of the methods couldn’t be assessed. The same dataset is used in Section 4.5 where the whole network of 45,101 genes is inferred and analysed. The dataset details are reported in Table 2.

**Table 2:** Characteristics of the real AD dataset

| GEO Accession # | Organism | Experiment Type | #Genes | #Time Points | Size of Network | Size of Matrix |
| --- | --- | --- | --- | --- | --- | --- |
| GSE32536 | *Mus musculus* | Expression profiling by array | 45101 | 16 | 2,034,100,201 | 721,616 |

#### 3.1.2. Synthetic Datasets

The Dialogue on Reverse Engineering Assessment and Methods (DREAM) [23] challenge is one of the de-facto standards for the assessment of biological network inference. Gene Net Weaver (GNW) [24] is a simulating tool that generates expression profiles for desired network size based on the DREAM networks. The synthetic datasets consisting of expression profiles along with their corresponding true networks were generated from GNW. The details of the synthetic datasets are reported in Table 3. The synthetic networks generated consist of up to 4000 genes (maximum genes possible in GNW is 4441) with sample points fixed at 210 and therefore expression profiles were of the maximum size of 840,000.

**Table 3:** Characteristics of the DREAM Challenge dataset used

| Dataset Name | Species | #Genes | #Interactions | #Time Points | Size of Matrix |
| --- | --- | --- | --- | --- | --- |
| E-1 | E. coli | 100 | 157 | 210 | 21000 |
| E-2 | E. coli | 500 | 1322 | 210 | 105000 |
| E-3 | E. coli | 1000 | 2019 | 210 | 210000 |
| Y-1 | Yeast | 2000 | 5256 | 210 | 420000 |
| Y-2 | Yeast | 3000 | 8029 | 210 | 630000 |
| Y-3 | Yeast | 4000 | 11323 | 210 | 840000 |

### 3.2. Candidate Inference methods

Out of the 15 different inference methods (discussed in section 2) considered in this work, three methods generated directed networks, while the rest inferred undirected networks. We used the R codes available online without any alterations for both the serial implementation as well as for the parallel implementation using our proposed framework. The default parameters were chosen wherever necessary for each method while executing them.

### 3.3. Inferring Large Regulatory Networks: A Novel Parallel Framework

The parallel computing paradigm has become indispensable in the inference of large scale GRN owing to the huge size of the expression profiles as well as inherent computational requirements for the existing GRN inference algorithms. In this regard, we envisaged putting forth a generic parallel framework that would be able to take in any inference algorithm and parallelly infer the network without the need to modify the inference algorithm. The non-requirement of any modification to the original inference algorithm and yet, the ability to parallelly infer the network is a unique feature of this framework. We adopted a simplistic mechanism to generate networks parallelly wherein the local or sub-networks are inferred concurrently using any particular inference method like plugins and subsequently all the sub-networks are merged to recreate the global network. Methodically, our approach is simple to implement yet effective.

The proposed framework works in five different phases. Phases 1 and 5 are executed in the serial environment while phases 2-4 are executed concurrently. The five phases of the framework (Figure 1) are elaborated as follows:

**Figure 1:**
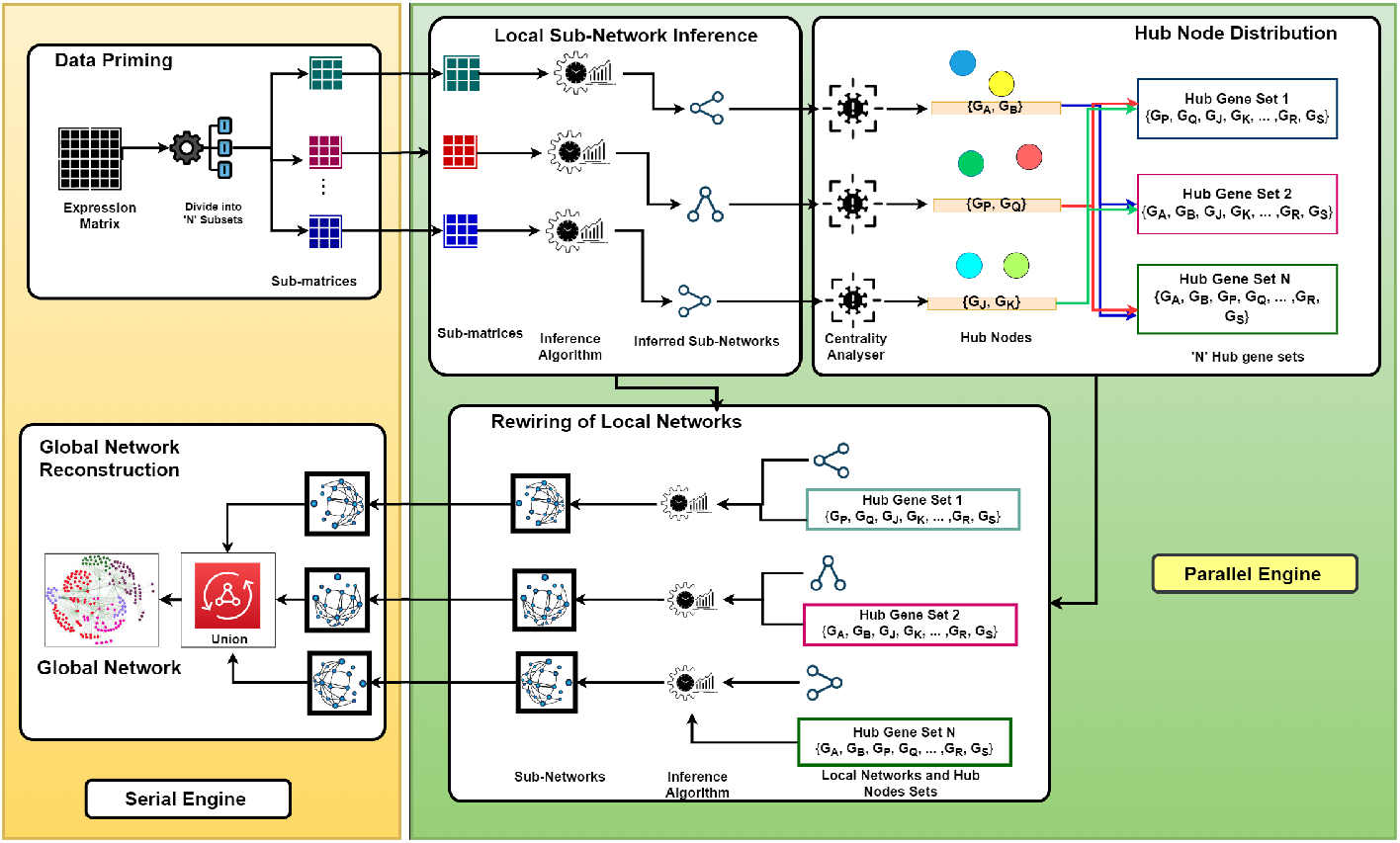
Workflow of the proposed generic parallel framework for inferring large scale network.

- **Data Priming:** The expression matrix consists of rows and columns where the rows indicate the expression profiles of all the genes while the columns depict the genes included in the matrix. Although this was the general format required by most algorithms, some changes (transposition) was needed in the format for some algorithms. In this phase, the input expression matrix is split into *p* subsets in order to be used concurrently in the succeeding phases. The expression matrix is subdivided horizontally in the following way. Given a matrix 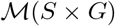, where *S* is the set of *n* expression profiles for gene *g_i_* ∈ *G* and *G* is the set of *m* genes (nodes), the partition can be as follows.

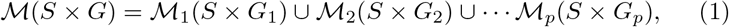

where, *G_i_* ⊂ *G* and 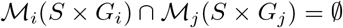.
- **Local Network Inference:** In this phase, local network or sub-networks are inferred from the split sub-matrices using any state-of-the-art inference method. It is noteworthy that the original implementation of the candidate algorithm is employed without altering any of its steps, treating it as a black box. The sub-networks are inferred parallelly in the different parallel execution units.
- **Hub Node Distribution:** The derived sub-networks from different units are disconnected as the expression matrices were initially divided into nonoverlapping subsets. Hence, a mechanism to re-wire the local sub-networks is required. The concept of *preferential attachment* [25, 1] inspired the relinking mechanism of the different sub-networks as real-world networks are usually scale-free. Preferential attachment means that the more connected a node (hub) is, the more likely it is to receive new links i.e. ”the rich-get-richer”. Nodes with a higher degree have a stronger ability to grab links added to the network. So it was perceived that the highest degree node or set of nodes or simply put the hub nodes, are most likely to form links with the genes from other sub-networks and hence can act as the bridge between the sub-networks. Therefore, in this phase, the hub genes in each sub-networks are determined and then the top two heavily connected genes in each sub-network are chosen for further sharing (Figure 1-Phase 3). The top two highly connected genes were chosen to tackle two possibilities. First, genes that are otherwise highly connected in the original network i.e., the actual hub genes, may not necessarily be the highly connected node in the sub-networks due to the possible loss of links during the division of the expression matrix. Second, the most highly connected gene of one sub-network may not be able to bridge it with other networks due to the possibility of not having any more additional links with the genes in the other sub-networks and therefore we use an additional hub gene to take no chances. However, the number of hub genes to be shared may be considered as a tuning parameter that depends on the data at hand. After the hub genes were identified, the *N* set of hub genes were shared to the *N* – 1 execution units, excluding the unit from where the set has been generated to avoid redundancy in the task of inference.
- **Rewiring Local Networks:** Herein, the sub-networks and the hub gene sets are employed to re-infer the network using the same inference algorithm chosen initially by the user to get the refined sub-network with overlapping nodes from different divisions. The parallel execution ends with this phase.
- **Global Network Reconstruction:** Now we have the final sub-networks wherein the links lost during expression matrix division have been reinstated thereby improving its quality. They are now integrated to output the final network. This amalgamation of sub-networks (or local adjacency matrices) gives the global network in the form of a global adjacency matrix.

### 3.4. Execution Platform

The R scripting language^2^ was used for all implementations and executions. We utilised an Intel(R) Core(TM) i5-8500, 3.00 GHz CPU based system with six (06) cores and 8 GB RAM that runs on Ubuntu 18.04.3 LTS and provides 6 concurrent threads of execution. The R implementation of the 15 candidate methods were obtained from *Bioconductor*^3^ and *CRAN* (The Comprehensive R Archive Network)^4^ and are mentioned in Table 1. The generic parallel framework too was developed in R scripting language (using *do Parallel*^5^ package) keeping in mind the availability of R implementations of the original methods. It helped easy coupling of the parallel platform with the candidate methods without re-coding or altering the original implementation.

### 3.5. Assessment Parameters

A network inference problem can be seen as a binary decision making task where the final decision about the possible existence (or absence) of an interaction between a pair of genes is nothing but a classification problem [1]. Therefore, the performance evaluation can be done with the metrics of machine learning like Area Under Receiver Operating Characteristic (AUROC) and Area Under Precision and Recall (AUPR) curves. ROC curves display the relative frequencies of true positives to false negatives for every predicted link of the edge list. Whereas the PR curves show the relative precision (the fraction of correct predictions) versus recall (also called sensitivity) that is equivalent to the true positive ratio [26]. The evaluation was focused on the GRN inference task and therefore only the most confident connections were taken into account. AUROC measures the capability of the algorithm in distinguishing edges, i.e., the higher the AUROC, the better the algorithm is capable of predicting an edge when it is present and not predicting it when it is absent. AUPR on the other hand measures the performance by evaluating the fraction of true edges among the edge predictions. It signifies the usefulness of the algorithm in comparison to others in identifying the actual edges. While the baseline for AUROC is fixed at 0.5 (0.5 meaning a bad classifier), the baseline of AUPR is determined by the ratio of positives and negatives.

The implementation of the generic parallel framework on candidate traditional inference methods was benchmarked against the serial implementation of the methods on the synthetic datasets using three parameters-execution time, AUROC and AUPR to gauge the enhancement brought about. The improvement in execution time was calculated using speedup. Speedup is the ratio of serial execution time to parallel execution time and is a metric to determine how fast the parallel execution is as compared to the serial execution of the same algorithm^6^.

## 4. Results and Discussion

In this section, we first report the scalability and performance assessment results of the different candidate GRN inference methods and then the performance of the proposed generic parallel framework in terms of speed-up achieved without compromising the qualitative performance of the original method. The candidate inference methods were also ranked based on their overall qualitative and quantitative performances. Lastly, we report the outcome of the inference and analysis of the GRN from the AD affected mouse model’s expression profiles using the best performing inference method.

### 4.1. Scalability analysis of the candidate GRN inference methods

The execution time trend of the original 15 inference methods for the datasets consisting of 100-10000 genes (nodes) is shown in Figure 2. The candidate methods were unable to infer larger networks of size 15000 nodes (genes) and above. As can be seen from the figure, C3MTC is the fastest method closely followed by RELNET, CLR and C3NET. G1DBN, GeneNet and MINE were the most expensive and showed exponential growth while inferring larger networks. Remarkably, G1DBN took 38 days to infer the network of 2000 genes. Due to the large time scale difference between G1DBN and the others, it could not be reported in the above mentioned plot. The trend depicted in the graph clearly shows that the execution time taken by the GRN methods increase drastically as the network size grows beyond 1000 genes. This drastic increase in the execution time demands an effective alternative in the form of parallel implementations so that the large-scale real networks can be inferred efficiently utilizing the above methods.

**Figure 2:**
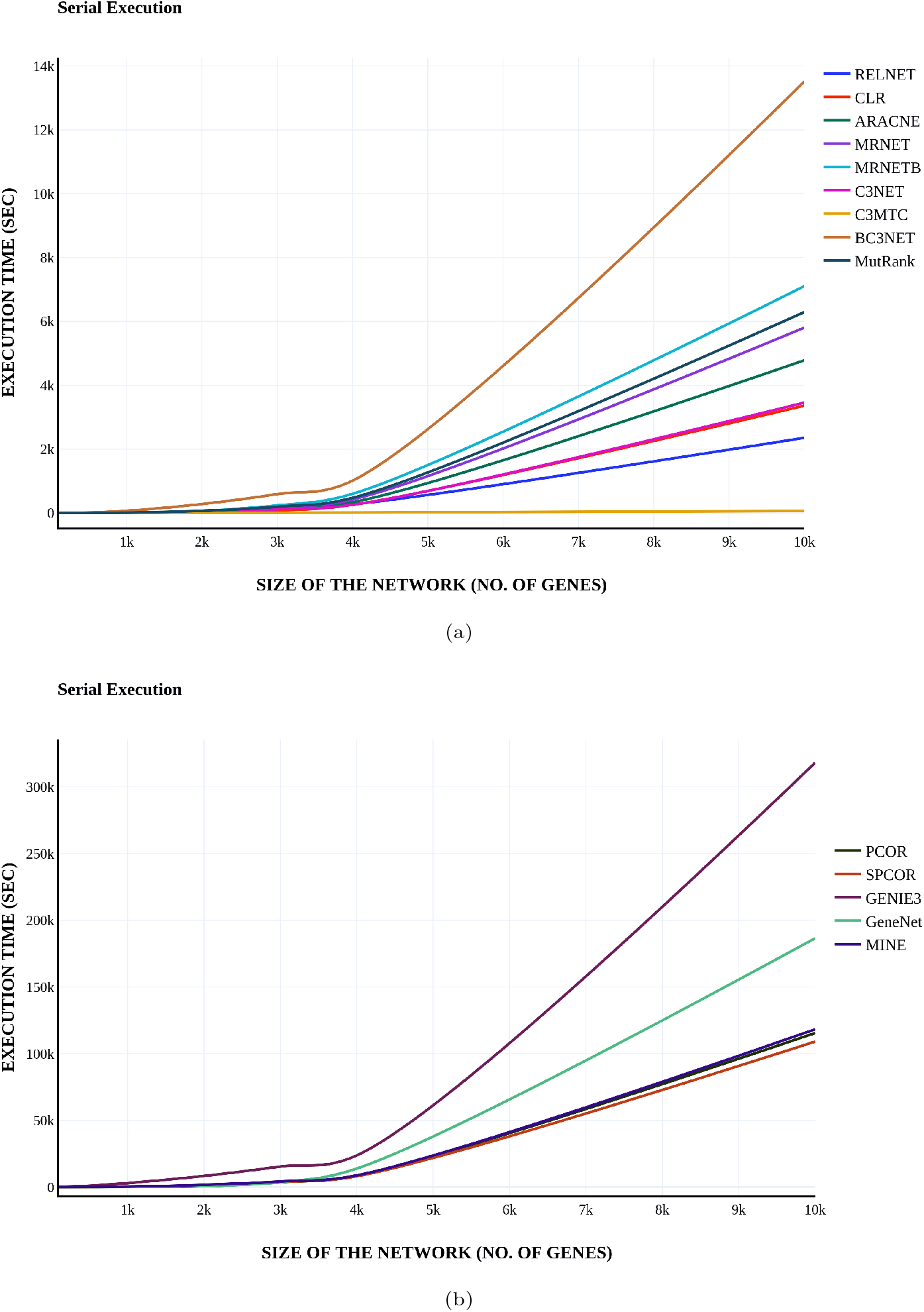
Execution time trend of serial execution of the GRN Inference Methods

### 4.2. Qualitative Assessment of the Serially Inferred Networks

The serially inferred networks were compared against the true networks (gold networks) for each of the synthetic datasets and the resultant AUROC (Area Under ROC) and AUPR (Area Under PR) values were graphically plotted as shown in Figure 3 and Figure 4. The results reveal that the GRN inference methods under consideration exhibit different behaviour across different datasets. It can safely be assumed that one particular method cannot be picked up as the best one for all datasets. For the network with 1000 genes, all methods show comparatively best performance as compared to the other networks. The performance of the methods drastically decreases with an increase in network size. On average, the highest AUROC obtained was only 0.601162. Although none of the methods obtains the best results throughout the datasets, as an overview G1DBN and CLR outperforms the other methods in their ability to distinguish between the presence or absence of edges between genes, closely followed by MRNETB and MRNET.

**Figure 3:**
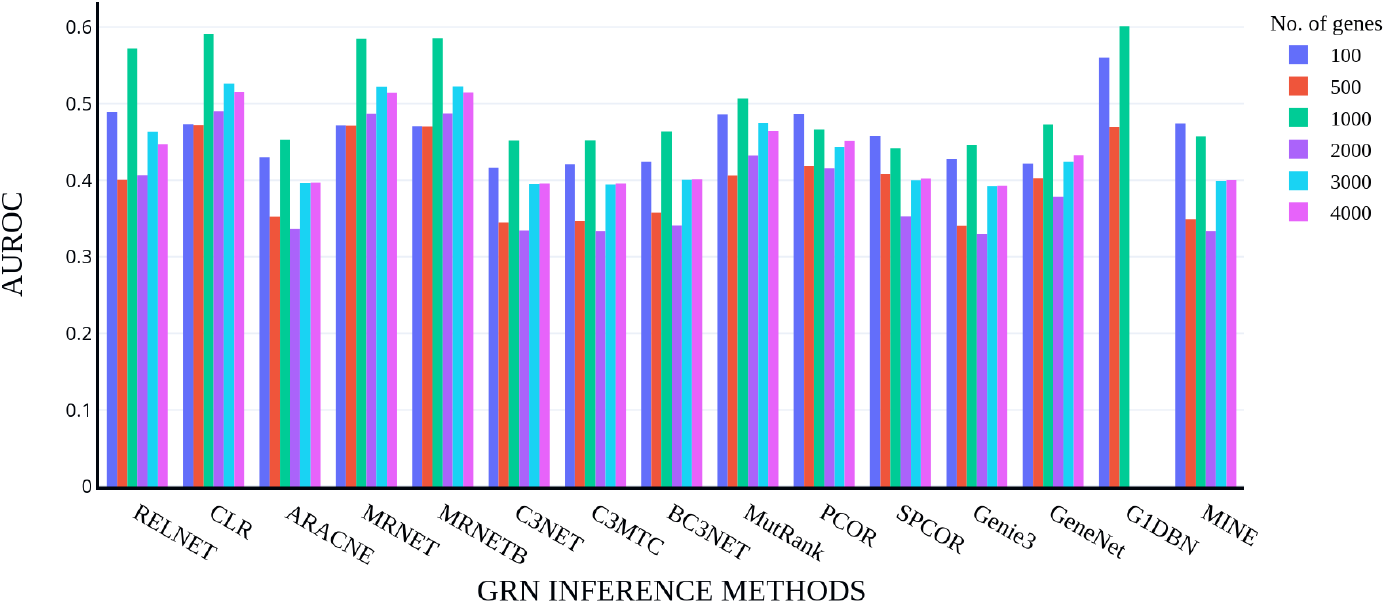
Comparison of the performance of the GRN Inference Methods

**Figure 4:**
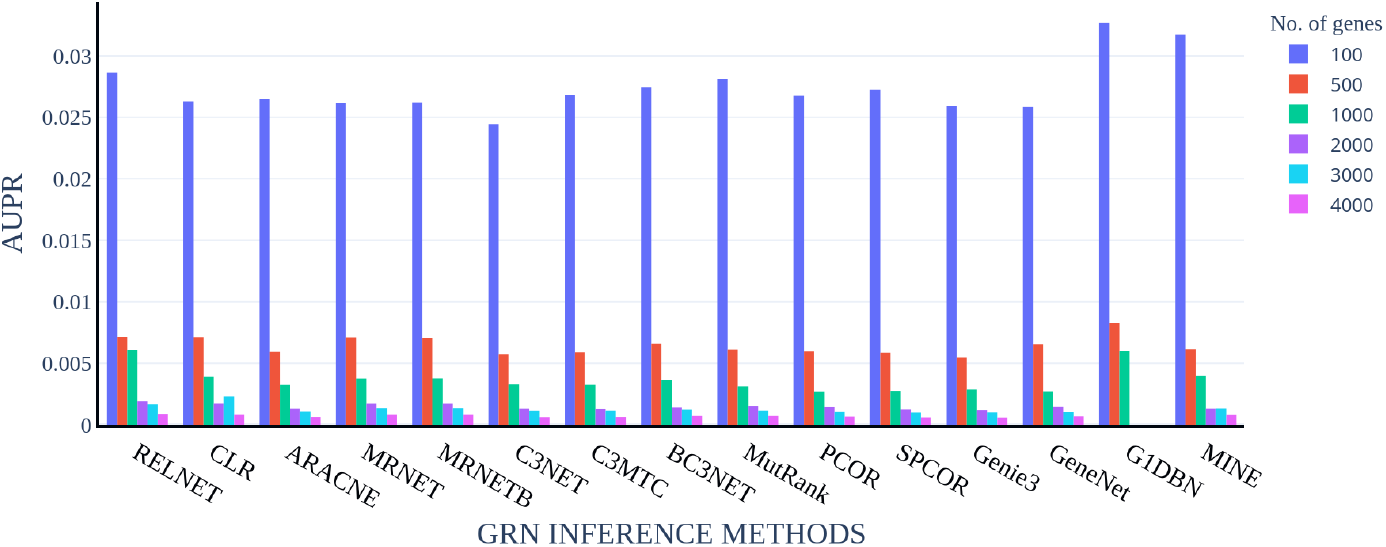
Comparison of the Prediction Accuracy of GRN Inference Methods

As far as the AUPR values are concerned, the baseline of the AUPR for the different datasets (100-4000) was determined as approximately 0.0157, 0.0053, 0.0021, 0.0013, 0.0009 and 0.0007 respectively. The low baseline can be attributed to the class imbalance since the number of edges is very few. Figure 4 shows that while all algorithms showcase a bare minimum AUPR score above the baseline, the performance decreases with the increase in the number of genes. As a general conclusion, it can be said that G1DBN, RELNET, MINE and CLR give the best AUPR scores and therefore are capable of predicting the edges much better than the other algorithms.

### 4.3. Ranking of candidate methods

The execution time, AUROC and AUPR score of each of the 15 inference algorithms were ranked separately on a scale of 1 to 15. The least time-taking algorithm was ranked first while the most time-exhaustive one was ranked last. On the other hand, the algorithm with the highest AUROC/AUPR score was ranked first while the one with the least was ranked the last. The combined average of all three ranks was then taken to finally get the overall ranking of all the methods. Therefore, the inference method that had the least average score was adjudged the best among the 15 inference methods. The ranking of the 15 inference methods based on execution time, AUROC score, AUPR score and overall performance is reported in Table 4. The overall ranking was necessary because some methods gave considerably good quality results but took ginormous time to execute. CLR emerged as the best performer in the overall ranking followed by RELNET, MRNET and G1DBN. It is to be noted that CLR is the only method to appear in the top 4 in all categories. This ranking table along with Table 1 can be useful for researchers to decide the trade-off between quality of inference and execution time.

**Table 4:** Ranking of the Inference Methods

| Rank | Based on Execution Time | Based on AUROC score | Based on AUPR score | Overall Ranking |
| --- | --- | --- | --- | --- |
| 1 | C3MTC | G1DBN | G1DBN | **CLR** |
| 2 | CLR | CLR | RELNET | **RELNET** |
| 3 | C3NET | MRNET | MINE | **MRNET** |
| 4 | RELNET | MRNETB | CLR | **G1DBN** |
| 5 | ARACNE | RELNET | BC3NET | **MRNETB** |
| 6 | MRNET | MutRank | MRNETB | **MutRank** |
| 7 | MutRank | PCOR | MRNET | **C3MTC** |
| 8 | MRNETB | GeneNet | MutRank | **BC3NET** |
| 9 | BC3NET | SPCOR | C3MTC | **MINE** |
| 10 | PCOR | MINE | SPCOR | **ARACNE** |
| 11 | SPCOR | BC3NET | ARACNE | **PCOR** |
| 12 | MINE | ARACNE | PCOR | **SPCOR** |
| 13 | GeneNet | C3MTC | GeneNet | **C3NET** |
| 14 | Genie3 | C3NET | Genie3 | **GeneNet** |
| 15 | G1DBN | Genie3 | C3NET | **Genie3** |

### 4.4. Effectiveness assessment of proposed generic parallel framework

In this section, we establish how the proposed generic parallel framework drastically improved the scalability of the candidate methods. We made the inference methods capable of inferring relatively larger networks on the same platform (where they failed earlier) without compromising on their original predictive quality.

First, we discuss the speedup achieved with the induction of the parallel platform that was calculated and has been reported in the Figure 5. The speedup achieved is different for each method for each dataset. For the methods that weren’t time-expensive, the speedup achieved is just fair initially, but as the size of the network increases, it becomes good enough. The major accomplishment of the parallel framework can be seen in the speedup achieved by the time-expensive methods. For example, the speed achieved by G1DBN went up to 729 meaning that the parallel execution was 729 times faster than the serial execution and similarly it went up to 313 for GeneNet, 158 for PCOR and 100 for MINE. Interestingly, candidate inference methods that were reported in Section 4.1 as incapable of inferring networks with more than 10000 genes were now inferring networks of even up to 20000 genes successfully. The crowning moment was when G1DBN that didn’t execute serially for more than 2000 genes executed for up to 10,000 genes. Then again, as the number of genes in the dataset increases, the speedup achieved also increases which shows that the parallel approach is scalable with the increase in the size of the datasets. It further established the superiority of our framework where even a simple PC with 4 CPU cores was able to infer large networks seamlessly.

**Figure 5:**
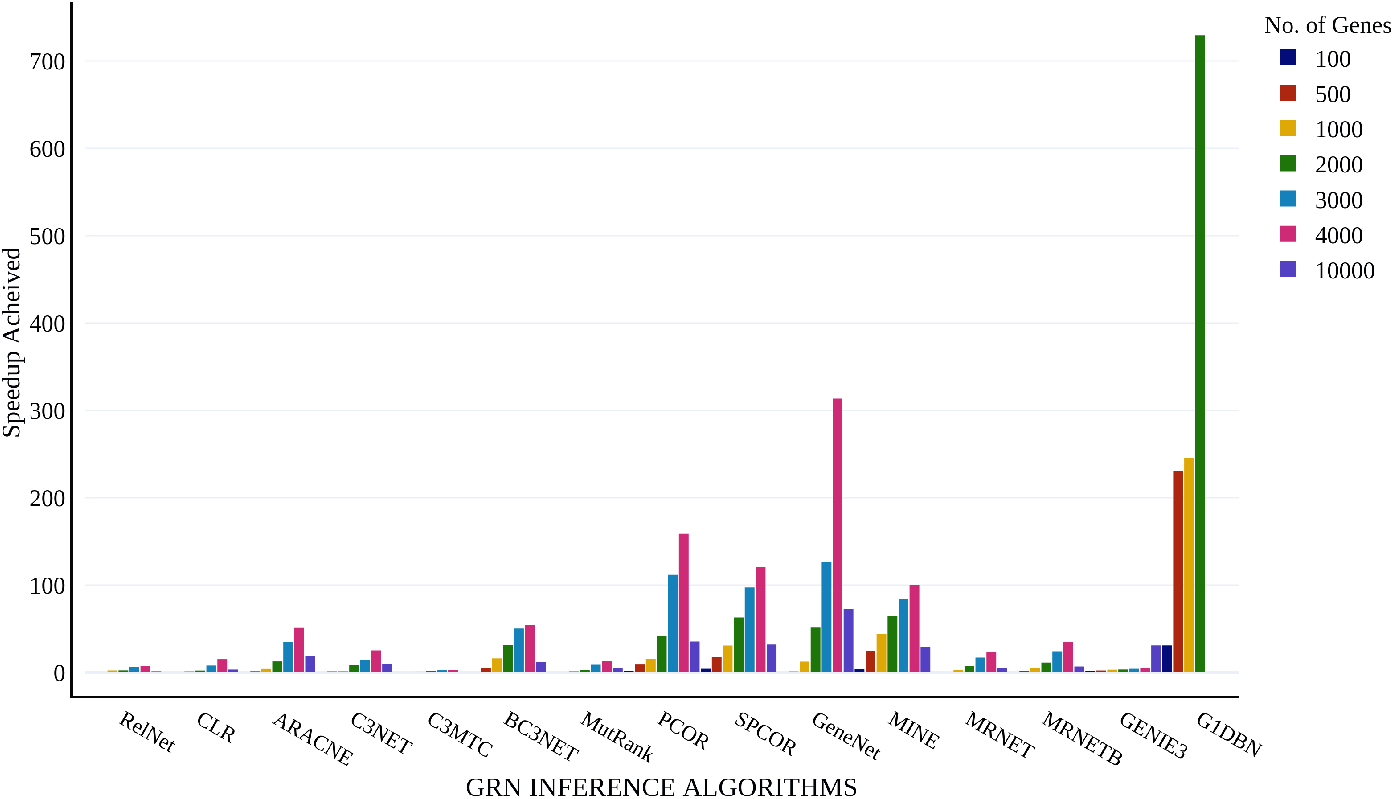
The Speedup achieved through parallel implementation of the 15 GRN Inference algorithms

The scalability of the parallel execution of the candidate methods using the generic parallel framework is depicted in Figure 6. Unlike what was seen in the scalability analysis of the serial execution of the methods, there is no steep rise as the number of genes increases herein. For example, in the case of CLR, the time for serial execution went from 0.02 to 251 secs when the size of the dataset increased from 100 to 4000 genes, while for the parallel execution it only went from 0.1 to 76 secs. G1DBN however could not be executed for more than 10000 genes (not shown in the graph due to large scale difference) due to R throwing an error upon trying for execution.

**Figure 6:**
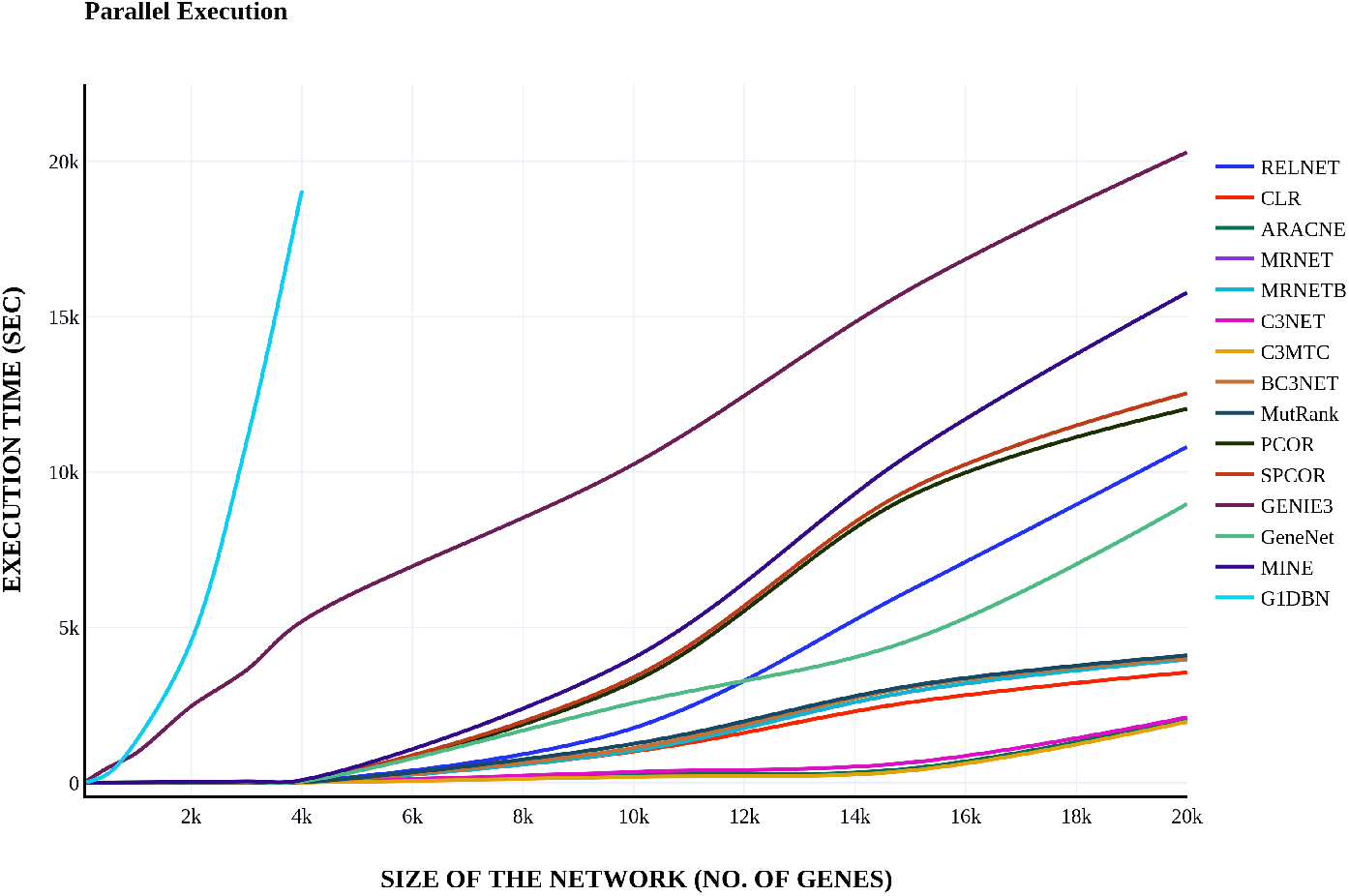
Execution time trend of the parallel implementation of the GRN Inference algorithms

The networks obtained from serial and parallel execution of each of the algorithms were compared against each other using AUROC and AUPR characteristics and the results are shown in Figure 7 and Figure 8. These networks were then again compared against the gold network and the results are depicted in Figure 9 and Figure 8. From all of these graphs, it can be safely concluded that the parallel approach proposed has been able to achieve what was envisaged in the beginning, i.e., inferring networks parallelly that will be as close as possible to the network otherwise obtained through serial execution. The generic parallel approach therefore is an opportunity for the research community to infer large scale networks without having to compromise on execution time or accuracy even if high computational resources are not available.

**Figure 7:**
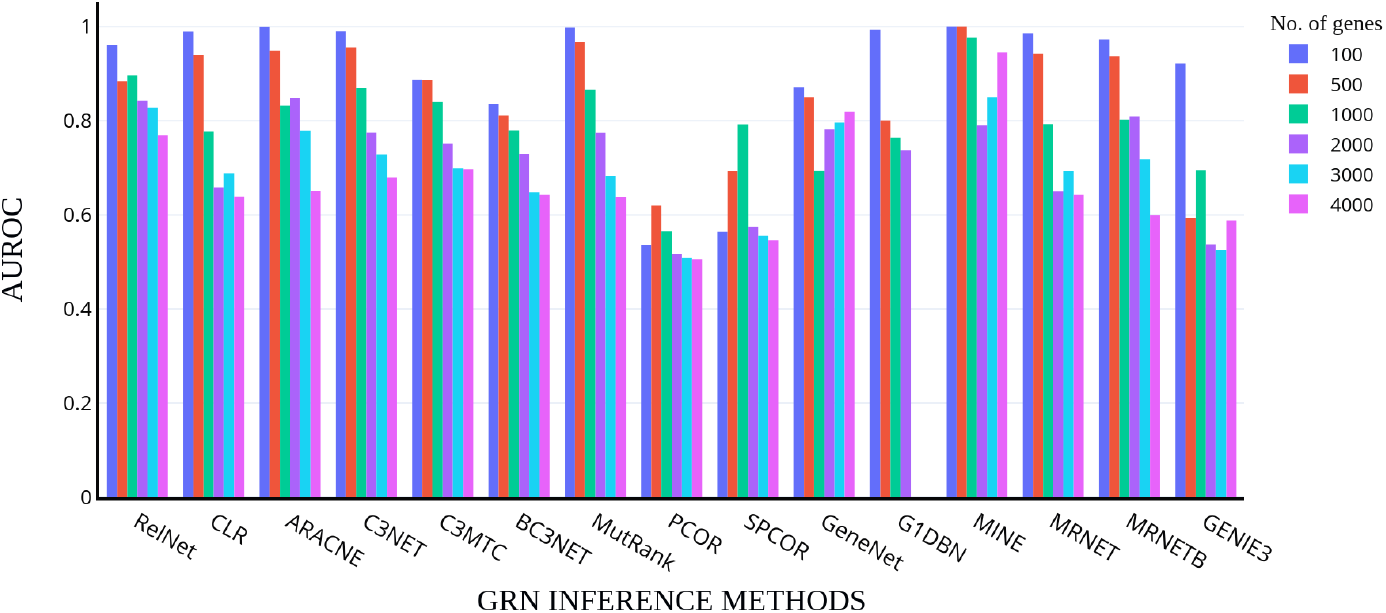
Comparison of the performance of the networks obtained from serial and parallel execution

**Figure 8:**
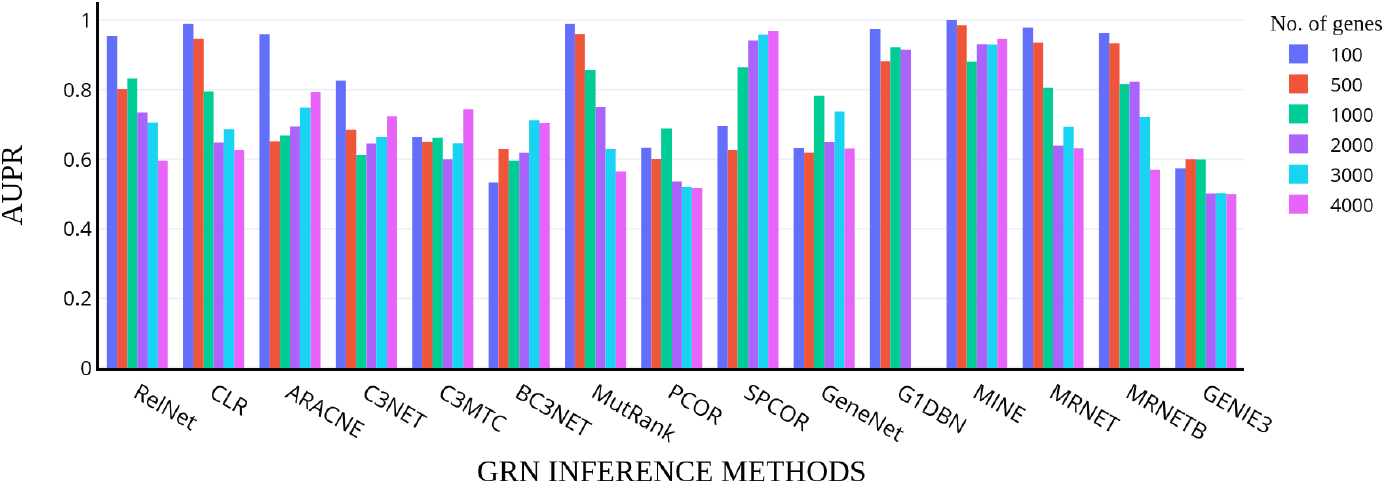
Comparison of the prediction accuracy of the networks obtained from serial and parallel execution

**Figure 9:**
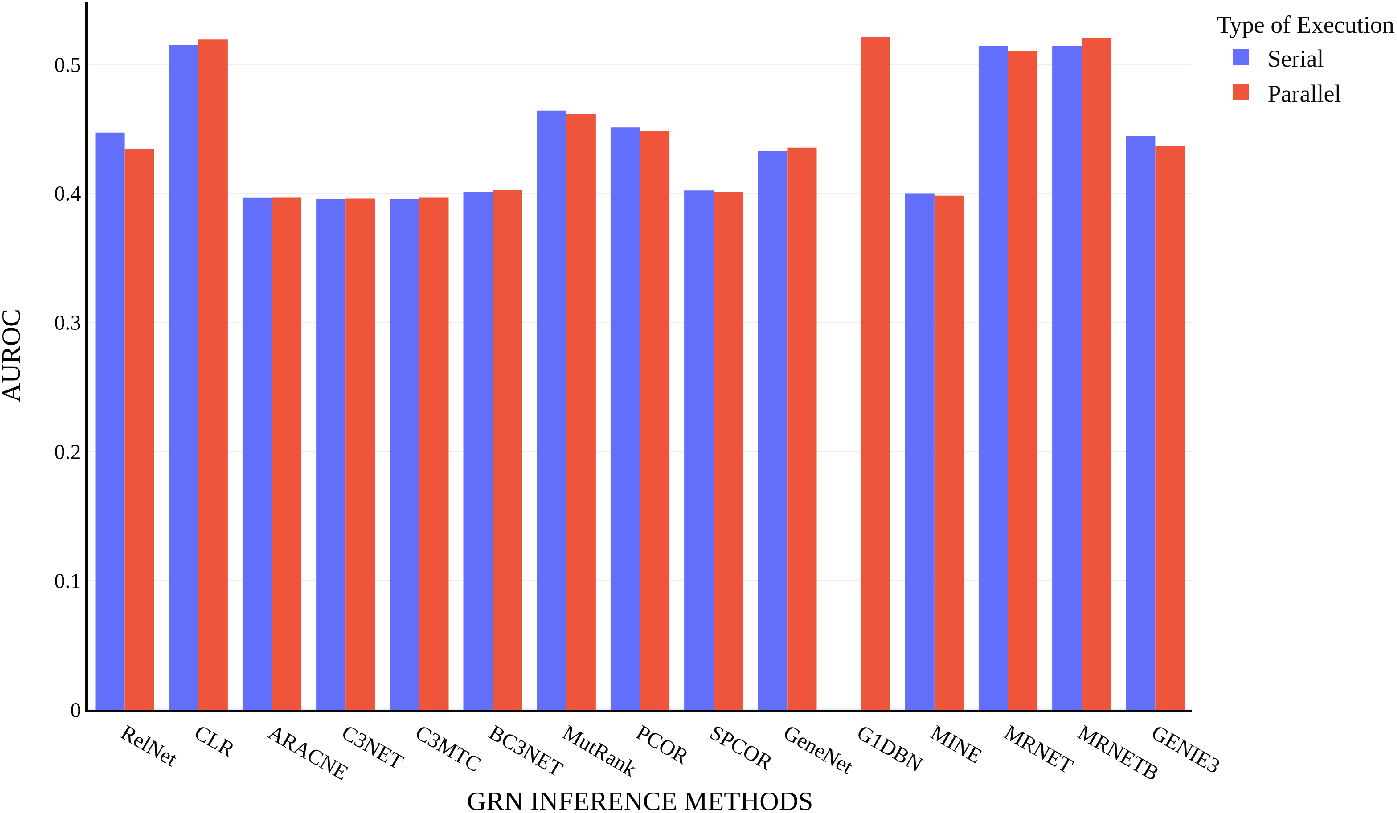
Comparison of the prediction accuracy of the networks obtained from serial and parallel execution against gold network for dataset with 4000 genes

**Figure 10:**
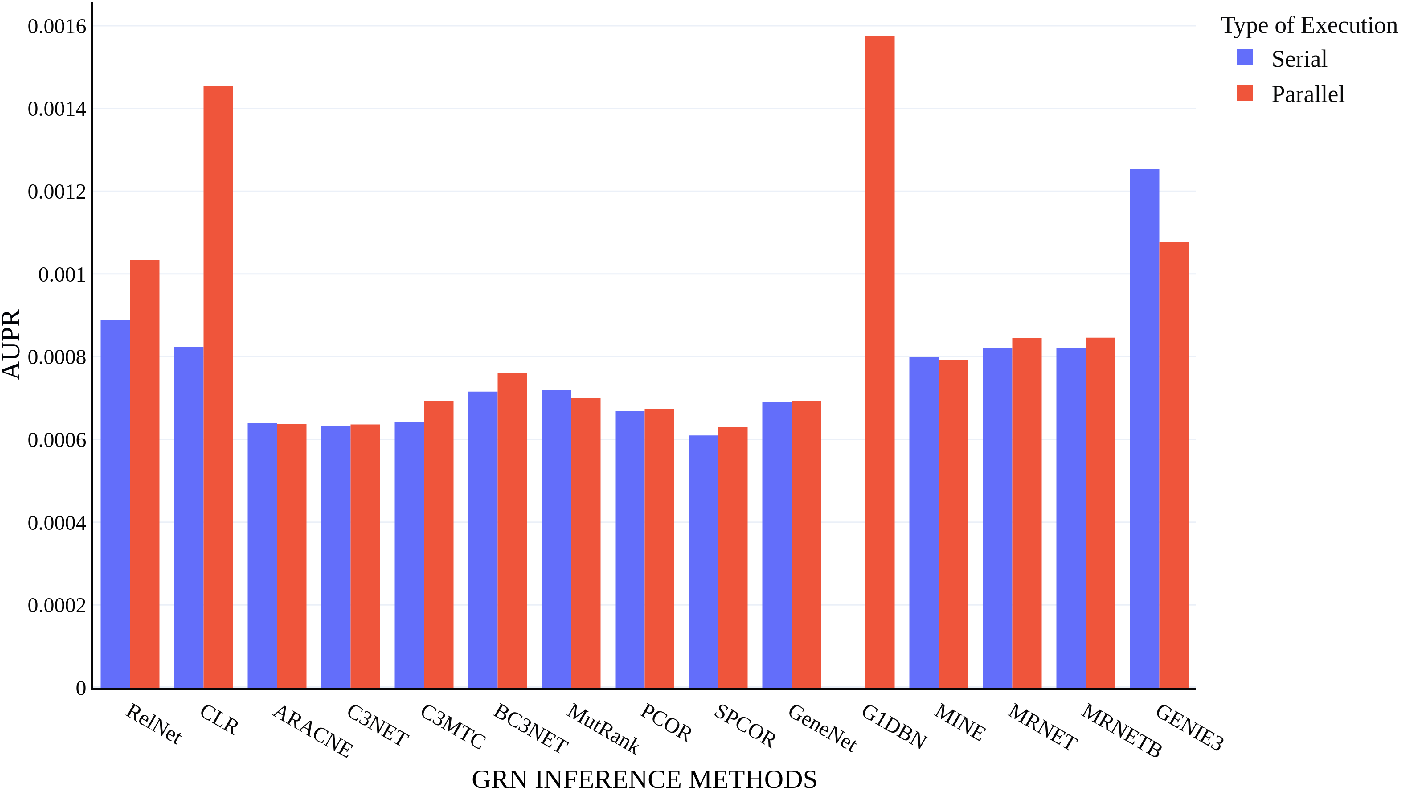
Comparison of the prediction accuracy of the networks obtained from serial and parallel execution against gold network for dataset with 4000 genes

For more detailed results on the execution time and prediction accuracy, please refer the Supplementary Material.

### 4.5. Inference and analysis of genome scale AD affected network

To demonstrate the applicability of the proposed parallel platform, we inferred the network of the real dataset chosen earlier that consisted of 45,101 genes and 16 samples using CLR which was the best performing algorithm. The large network couldn’t be inferred in our previously specified environment because of insufficient RAM. Therefore, we used a workstation with a higher configuration to run the parallel CLR. The workstation was built of dual 2.1GHz Intel processors, 16 cores and 256 GB RAM. Even in the above workstation, none of the serial methods was able to handle the large network of 45,101 genes. The parallel CLR took almost 1.83 hrs to complete the execution. The resultant network was further analysed in order to obtain the network characteristics.

First, we calculated the degree distribution of the inferred network and plotted it as depicted in Figure 11. The figure shows that while most of the nodes have a relatively small degree, quite a few nodes have a very large degree, indicating the presence of few hub nodes. These hub nodes have caused the network’s degree distribution to have a long tail. From the figure it is apparent that the inferred network loosely follows scale-free properties with power-law like degree distribution [25, 1]. It is worth mentioning that true scale-free networks are rare in reality [27].

**Figure 11:**
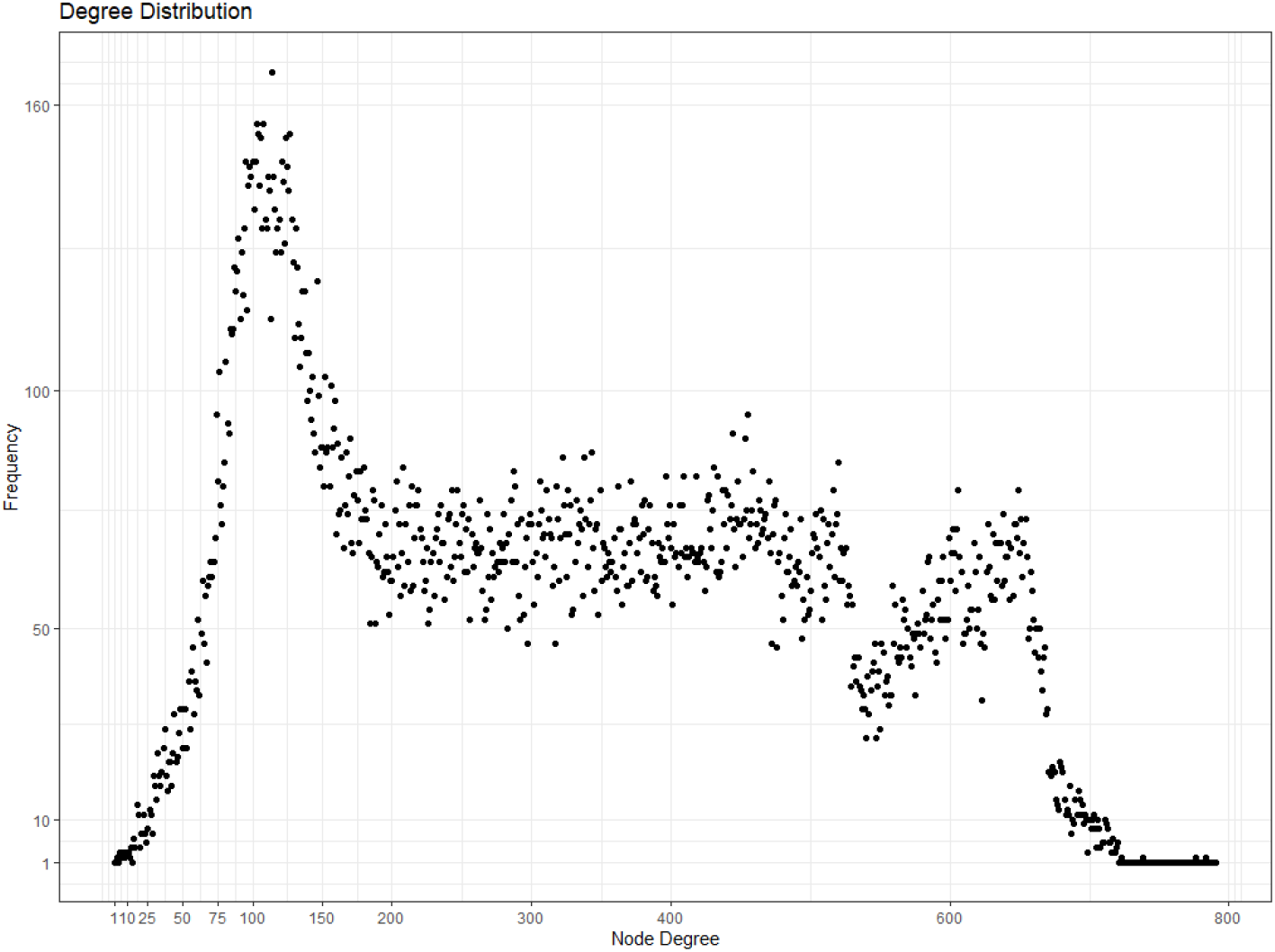
Plot of the degree distribution of the inferred genome scale AD affected GRN

Owing to our proposed framework, CLR was capable of inferring a large network for the first time from genome scale expression profiles, which was previously unknown due to almost no experiments done on large datasets. The network characteristics, that is, density, average path length, clustering coefficient along with the global topological centralities of the inferred network were quantified next. The inferred network has a low density of 0.0129563 because out of the 45101 × 45101 (2,034,100,201) edges it could have had, only 1,048,576 are present. The average path length of 2.0896876 suggests that the genes are interconnected through a very short path. The low global clustering coefficient score of 0.1498094 shows the strong influence of the hub nodes in the network. The global degree centrality of the inferred network is neither high nor low score of 0.6733727 because although the network is distributed, some hub genes are interconnected. This again is the reason why the global closenness centrality score is neither high nor low score of 0.560417. The low global betweenness centrality score of 0.0340206 is probably because of the multiple paths that might exist since the network is spread out. The global Eigenvector centrality is a high score of 0.95852 because the hub genes are highly connected among themselves.

### 4.6. Identification and Analysis of Hub Genes in the genome scale AD affected network

Various researches reported the significance of central or hub genes in a network and their possible role in causing diseases [28, 29, 3, 30, 31]. The node centrality was computed with the help of the four popular centrality measures, namely, degree, betweenness, closeness and eigenvector centrality. The topological centrality scores of each gene were then used to rank the genes. However, since different centrality measures were found to vary in their scores for the same gene, we consolidated the scores using a simple Rank Product (RP) approach [32]. Given a vector of ranking scores for each gene *i,* {*R*_1_, *R*_2_, ⋯, *R_n_*} the *RP* of *i* can be calculated as follows.

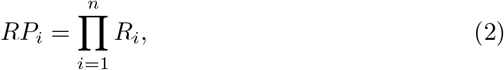

which is the product of *n* centrality scores for the node/gene *i*.

The top 5% of the ranked genes were then deemed as the key genes in this dataset. A total of 2255 genes were thus identified. The top 15 genes and their corresponding RP score along with their topological centrality scores are presented in Table 5. The complete table can be found in the Supplementary Material. We then proceeded to analyse the interactions among the top 1% genes by extracting a sub-network closely related to those genes alone and then visualising the sub-network as shown in Figure 12. The visualisation helped us find out that with the exception of two hub genes, the rest 449 genes formed a highly interacting sub-network implying the presence of ”rich-club” in the network and also hints that the key genes partake in similar biological processes together and hence could be potential biomarkers.

**Figure 12:**
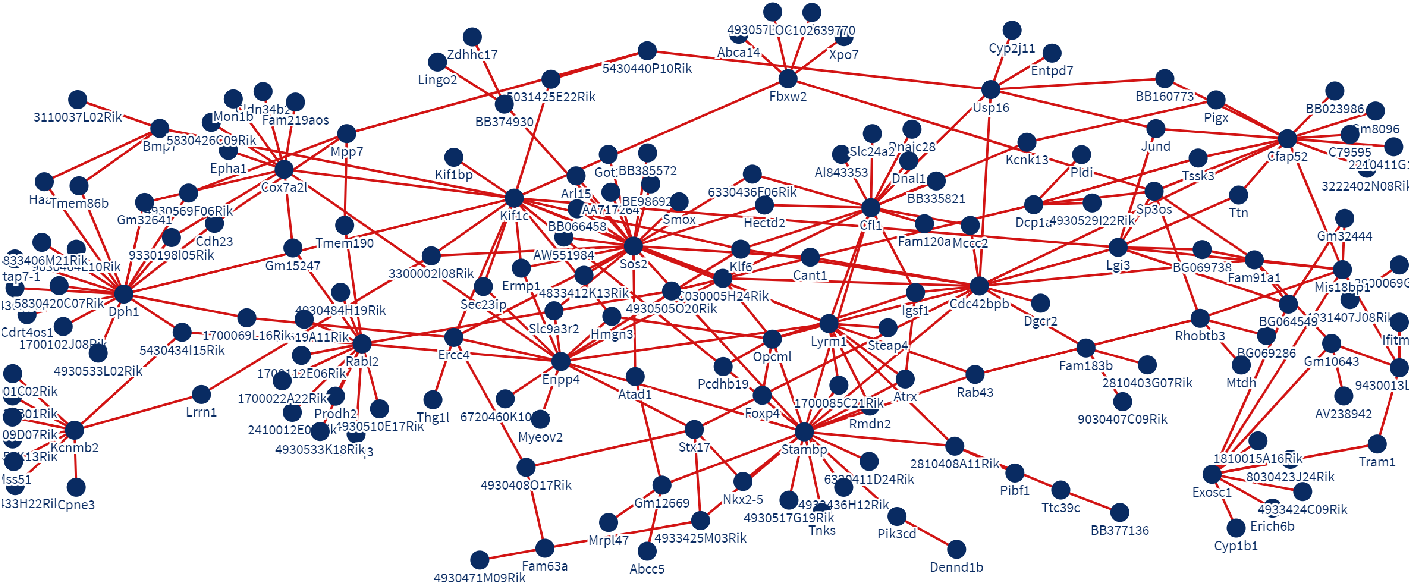
Visualisation of the “rich club” sub-network formed by the top 1% key genes extracted from the inferred AD affected GRN

**Table 5:**
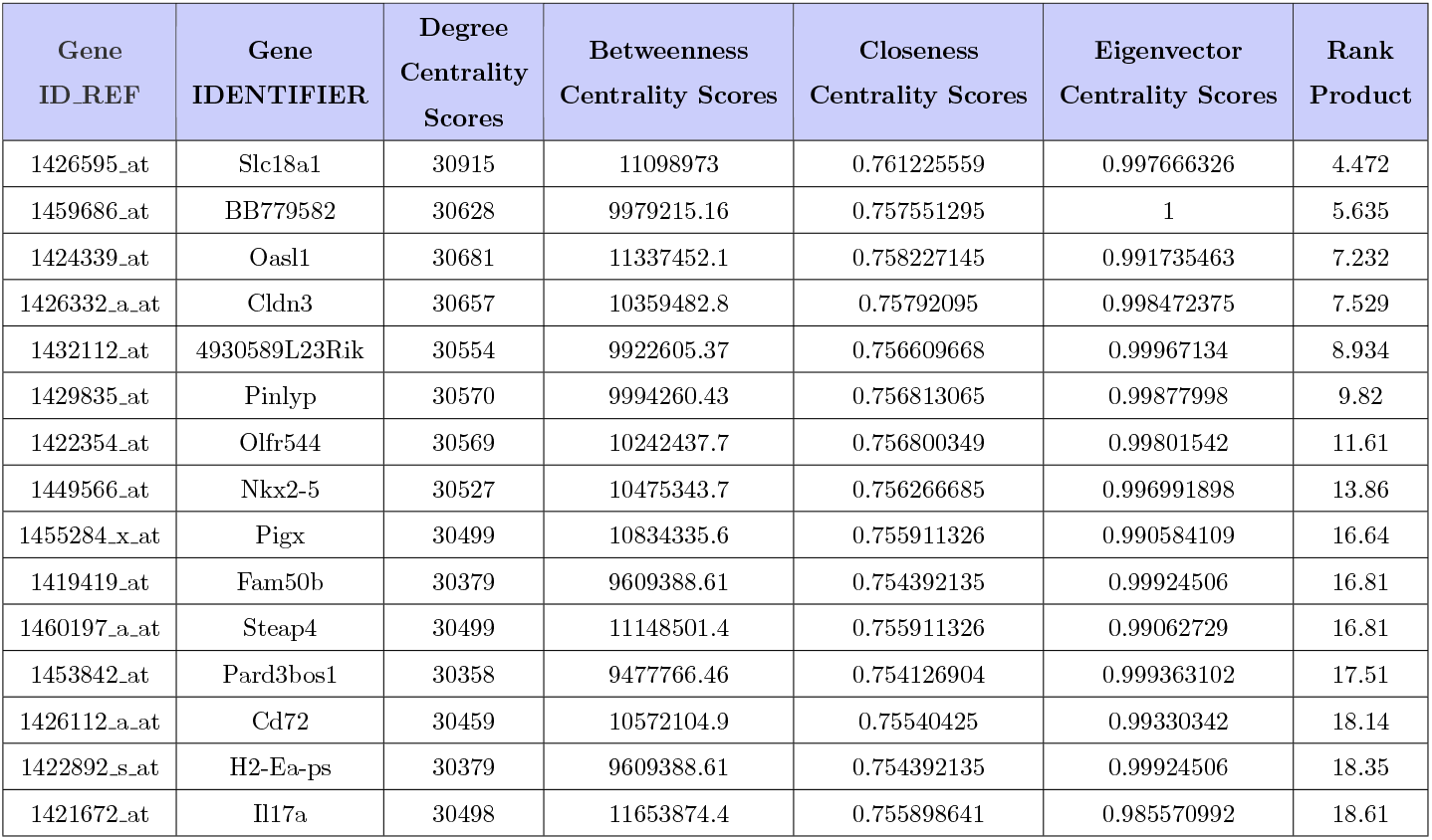
Centrality scores of the top 15 significant genes obtained (in order of their rank) using RP algorithm

Then the important question was how to validate if they are indeed the key genes or not? For this purpose, the significant genes (1936) reported in the work of [22], was compared with the key genes that were ranked in this work and it was found that more than 90% (1839) of genes matched. When the top 8% genes (3608) were taken for comparison, all the 1936 genes could be detected. This finding not only revealed the combined ability of CLR and our generic parallel framework to infer the large network efficiently but also pointed out the possibility of the presence of more significant genes than that was reported earlier.

Gene analysis was then performed using DAVID [33], GeneCards [34] and MalaCards [35] on the top 50 genes to verify our finding that the key genes probably partake in similar biological processes due to which they are highly inter-connected thus forming a ”rich club”. The results of the gene analysis of the top 15 genes are reported in Table6 while the complete gene analysis of the 50 top hub genes can be found in the Supplementary Material. We first searched the DAVID database using the gene IDs to get the corresponding DAVID gene name which was further used to query the gene category, the GeneCards Inferred Functionality Scores (GIFts) of the genes, disorders associated with the genes and their corresponding association scores from GeneCards and the type of disorders from MalaCards. Out of the 50 genes analysed, 29 were various disease-causing genes with high association scores, which included mental diseases, cardiovascular diseases, reproductive diseases, neuronal diseases and cancer diseases to name some while the others were either pseudogenes or genes exclusive to mouse. The question that how come the genes that cause different diseases are highly interconnected naturally arose. It is important to first mention that Searcy et.al. [22] in their work used PIO to treat the AD affected mice and reported that the several novel targets of PIO were found associated with cognition as well as estrogenic and glutamatergic pathways and that PIO also targeted processes like lipid/fatty acid metabolism, mitochondrial energy processes and synaptic neuro-transmission. The drug PIO, therefore, is the first reason that the divergent disease-causing genes were found interconnected. The second reason is that most of the identified hub genes are not limited to just one type of disease. This ability of the genes to cause multiple diseases resulted in the high level of interactions among them. The 1936 significant genes that were reported played a role in inflammatory processes, glutamatergic neurotransmission, lipid metabolism, cholesterol transport and female hormone processes and so does the top 50 hub genes identified in this paper. This shows that the network inferred by the parallel CLR was effective in pointing out the disease-causing genes. Therefore, the generic parallel framework has been established as efficient to infer large scale networks. This achievement opens up the opportunity for the research community to infer large scale networks with any method of inference to obtain the type of network that they want and further process it without having to go about with alternatives.

**Table 6:**
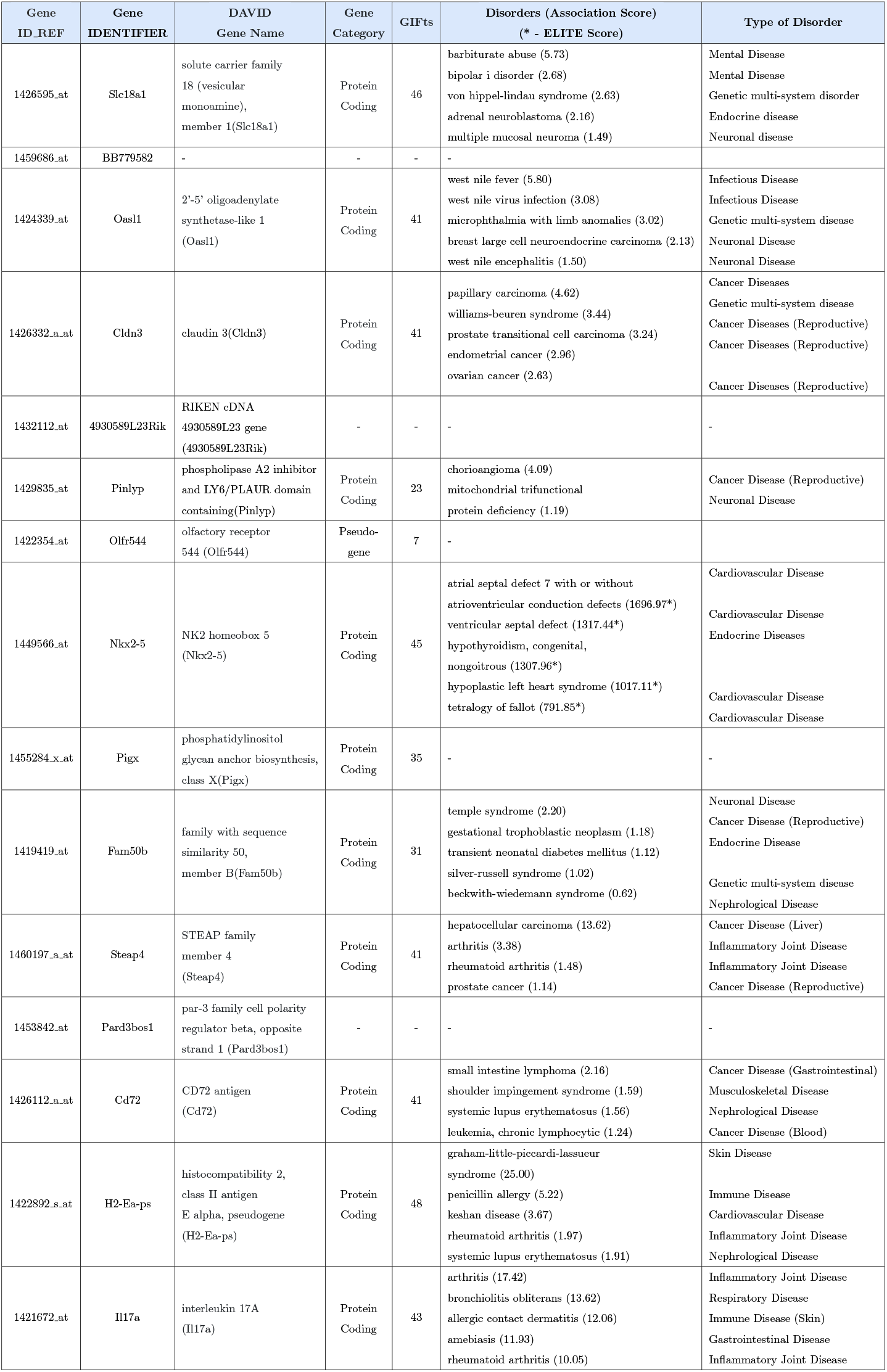
Gene Analysis of the top 15 genes in the inferred AD affected GRN

## 5. Conclusion

The inference of Gene Regulatory Networks (GRNs) continues to remain an important open challenge in computational biology. The goal of inference is to obtain the sparse topological structure and the parameters that help quantitatively understand and reproduce the dynamics of a biological system based on time-series of gene expression data. Obtaining the gene regulatory networks is not GRN inference’s raison d’être; the application of the inferred networks to help solve several different biological and biomedical problems and find treatments or prevention mechanisms for dreaded genetic disorders is why a lot of time, effort and resources are pooled in the area of computational biology. This paper, therefore, embarked on an idea to present before the research community a generic parallel inference framework using which any original inference algorithm without any alterations, can parallelly run on humongous datasets in the multiple cores of the CPU to provide efficacious inferred networks. Not only did we take up 15 inference methods and assess their scalability and performance but we also implemented the same methods using the proposed framework and showed how successful it was in overcoming the scalability limitations without compromising on the quality of the network inferred. We further inferred a real network from a humongous expression profile consisting of 45,101 genes and further analysed it to prove the efficacy of the generic parallel framework. The framework will prove pivotal in reviving the use of those inference methods that were deemed incompetent of inferring large networks earlier. The inference of GRN itself can be a tough task for researchers from non-computational backgrounds and therefore using the generic parallel framework might only add to the trouble. Hence we are now looking forward to developing a good interface for executing the generic parallel framework on any inference methods at the click of a button. This will not only ease the process of inference but also extend the ability to infer networks to virtually anyone. We also intend to include multi-source data for inference since micro-array data can be inconclusive at times. In the future, we also look forward to executing the framework in GPU to further lessen the execution time required.

## Acknowledgement

We extend our sincere thanks to Sk. Atahar Ali for his help in coding the preliminary version of the proposed framework and also to Mr. Sumit Dutta of NetRA Lab for his technical support in implementing the framework.

## Footnotes

1 https://www.ncbi.nlm.nih.gov/

2 https://www.r-project.org

3 https://www.bioconductor.org

4 https://cran.r-project.org

5 https://cran.r-project.org/web/packages/doParallel/index.html

6 https://software.intel.com/content/www/us/en/develop/articles/predicting-and-measuring-parallel-performance.html

